# Tumor cell-adipocyte gap junctions activate lipolysis and contribute to breast tumorigenesis

**DOI:** 10.1101/277939

**Authors:** Jeremy Williams, Roman Camarda, Serghei Malkov, Lisa J. Zimmerman, Suzanne Manning, Dvir Aran, Andrew Beardsley, Daniel Van de Mark, Rachel Nakagawa, Yong Chen, Charles Berdan, Sharon M. Louie, Celine Mahieu, Daphne Superville, Juliane Winkler, Elizabeth Willey, John D. Gagnon, Seda Kilinc Avsaroglu, Kosaku Shinoda, Matthew Gruner, Hiroshi Nishida, K. Mark Ansel, Zena Werb, Daniel K. Nomura, Shingo Kajimura, Atul J. Butte, Melinda E. Sanders, Daniel C. Liebler, Hope Rugo, Gregor Krings, John A. Shepherd, Andrei Goga

**Affiliations:** Department of Cell & Tissue Biology, University of California, San Francisco, San Francisco, CA, USA; Biomedical Sciences Graduate Program, University of California, San Francisco, San Francisco, CA, USA; Department of Radiology & Biomedical Imaging, University of California, San Francisco, San Francisco, CA, USA; Department of Biochemistry, Vanderbilt University School of Medicine, Nashville, TN, USA; Jim Ayers Institute for Precancer Detection and Diagnosis, Vanderbilt-Ingram Cancer Center, Nashville, TN, USA; Department of Pathology, Vanderbilt University School of Medicine, Nashville, TN, USA; Institute for Computational Health Sciences, University of California, San Francisco, San Francisco, CA, USA; Department of Medicine, University of California, San Francisco, San Francisco, CA, USA; Diabetes Center, University of California, San Francisco, San Francisco, CA, USA; Eli and Edythe Broad Center of Regeneration Medicine and Stem Cell Research, University of California, San Francisco, San Francisco, CA, USA; Department of Chemistry, University of California, Berkeley, Berkeley, CA, USA; Department of Molecular & Cell Biology, University of California, Berkeley, Berkeley, CA, USA; Department of Nutritional Sciences & Toxicology, University of California, Berkeley, Berkeley, CA, USA; Department of Anatomy, University of California, San Francisco, San Francisco, CA, USA; Helen Diller Family Comprehensive Cancer Center, University of California, San Francisco, San Francisco, CA, USA; Department of Microbiology & Immunology, University of California, San Francisco, San Francisco, CA, USA; Sandler Asthma Basic Research Center, University of California, San Francisco, San Francisco, CA, USA; Division of Endocrinology, Diabetes and Metabolism, Beth Israel Deaconess Medical Center, Harvard Medical School, Boston, MA, USA; Howard Hughes Medical Institute, Chevy Chase, MD USA; Department of Pathology, University of California, San Francisco, San Francisco, CA, USA; Faculty of Biology, Technion, Israel Institute of Technology, Haifa, Israel; The Taub Faculty of Computer Science, Technion, Israel Institute of Technology, Haifa, Israel; Tongji Medical College, Huazhong University of Science and Technology, Wuhan, China; Center for Cancer Research, Medical University of Vienna, Austria; Department of Medicine and Molecular Pharmacology, Albert Einstein College of Medicine, Bronx, NY, USA; Department of Radiation Oncology, Perelman School of Medicine, University of Pennsylvania, Philadelphia, PA, USA; Cancer Center, University of Hawaii, Honolulu, HI, USA

**Keywords:** Breast cancer, triple-negative breast cancer, TNBC, adipocyte, gap junction, lipolysis, cAMP, connexin 31, Cx31, GJB3

## Abstract

A pro-tumorigenic role for adipocytes has been identified in breast cancer, and reliance on fatty acid catabolism found in aggressive tumors. The molecular mechanisms by which tumor cells coopt neighboring adipocytes, however, remain elusive. Here, we describe a direct interaction linking tumorigenesis to adjacent adipocytes. We examine breast tumors and their normal adjacent tissue from several patient cohorts, patient-derived xenografts and mouse models, and find that lipolysis and lipolytic signaling are activated in neighboring adipose tissue. We find that functional gap junctions form between breast cancer cells and adipocytes. As a result, cAMP is transferred from breast cancer cells to adipocytes and activates lipolysis in a gap junction-dependent manner. We identify connexin 31 (GJB3), which promotes receptor triple negative breast cancer growth and activation of lipolysis *in vivo*. Thus, direct tumor cell-adipocyte interaction contributes to tumorigenesis and may serve as a new therapeutic target in breast cancer.

**One sentence summary:** Gap junctions between breast cancer cells and adipocytes transfer cAMP and activate lipolysis in the breast tumor microenvironment to support growth.

## Materials and Methods

### 3CB patient population

Five hundred women with suspicious mammography findings (BIRADS 4 or greater) were recruited and imaged before their biopsies using a 3-compartment decomposition dual-energy mammography protocol (3CB). This was multicenter study with two recruitment sites: University of California at San Francisco and Moffitt Cancer Center, Tampa, Florida. All patients received a biopsy of the suspicious area, and breast biopsies were clinically reviewed by the pathologists. A subset of pathology proven triplenegative (*n* = 6) and receptor-positive (*n* = 40) invasive cancers were selected for this study. All women received both craniocaudal (CC) and mediolateral-oblique (MLO) views. Exclusion criteria for the study were no prior cancer, biopsies, or breast ipsilateral alterations, and no occult findings.

### 3CB imaging protocol

The 3CB method combines the dual-energy X-ray mammography attenuations and breast thickness map to solve for the three unknowns water, lipid, and protein content [15]. We used Hologic Selenia full-field digital mammography system (Hologic, Inc.) to image women with 3CB. Two dual energy mammograms were acquired on each woman’s affected breast using a single compression. The first exposure was made under conditions of regular clinical screening mammogram. The second mammogram was acquired at a fixed voltage (39 kVp) and mAs for all participants. A high energy exposure (39 kVp/Rh filter) was made using an additional 3-mm plate of aluminum in the beam to increase the average energy of the high energy image. We limited the total dose of this procedure to be approximately 110% of the mean-glandular dose of an average screening mammogram. The images were collected under an investigational review board approval to measure breast composition. The breast thickness map was modeled using the SXA phantom [40](*34*). The thickness validation procedure concluded in a weekly scanning of specially designed quality assurance phantom [41]. The calibration standards and 3CB algorithms are described in full elsewhere [15, 42]. The region of interests of lesions and three surrounding rings of 2 mm distance outward from lesion boundary were derived for water, lipid, and protein maps. The median lipid measures of regions of interest within lesions, three rings outside of lesions, differences and ratios between lesions and rings were generated for both CC and MLO mammograms. Average values of generated variables of two views were used.

### Histological sectioning, hematoxylin and eosin staining, and adipocyte area quantification

Invasive breast carcinomas were obtained from the Pathology Departments of the University of California San Francisco (San Francisco, CA) and Moffitt Cancer Center (Tampa, FL). The study population included 39 hormone receptor positive tumors (32 ER positive (+)/PR+/HER2 negative, 2 ER+/PR-/HER2-, 4 ER+/PR+/HER2+, and 1 ER+/PR-/HER2+), 6 triple negative (ER-/PR-/HER2-) tumors, and 1 ER-/PR-/HER2+ tumor. Thirty-nine tumors were invasive ductal carcinomas and 7 were invasive lobular carcinomas. Tissue was fixed in 10% formalin and embedded in paraffin, and 4 micron sections were cut for hematoxylin and eosin (H&E) and immunohistochemical ER, PR, and HER2 staining, as well as HER2 fluorescence *in situ* hybridization (FISH) for a subset of tumors. ER, PR, and HER2 were scored according to ASCO/CAP guidelines [43, 44]. An H&E-stained slide demonstrating tumor and sufficient (at least 0.5 cm) NAT was chosen from each of 11 tumors with available slides and subjected to whole slide scanning at 400× magnification using an Aperio XT scanner (Leica Biopsystems, Buffalo Grove, IL). Images were visualized using ImageScope software (Leica Biosystems). For each tumor, 4 representative images at 50X magnification (at least 50 adipocytes per image) from R1 and R3 were analyzed using Fiji imaging software with the opensource Adiposoft v1.13 plugin [45]. This study was approved by the institutional review board of the respective institutions.

### cAMP-dependent lipolysis signature

The cAMP-dependent lipolysis gene signature was generated using RNA-seq data of cAMP-treated adipocytes [18]. Differentially expressed genes were sorted according to their *P* value and the top 100 upregulated genes were chosen for the signature. This signature was then used to calculated enrichment scores using the single-set gene set enrichment analysis (ssGSEA) method [20]. “cAMP 100 signature” enrichment scores were calculated for a dataset containing multiple samples from multiple regions surrounding breast tumors [19]. The dataset includes samples from the tumor itself (*n* = 9), and NAT 1 cm (*n* = 7), 2 cm (*n* = 5), 3 cm (*n* = 3) and 4 cm (*n* = 4) away from the tumor, in addition to healthy normal samples (*n* = 10). The spatial data set of multiple regions surrounding breast tumors was download from EMBL-EBI ArrayExpress (Accession E-TABM-276). Raw CEL files were downloaded and processed using custom Affymetrix GeneChip Human Genome U133 Plus 2.0 CDF obtained from BrainArray [46]. The processing and normalization were performed using the Robust Multi-array Average (RMA) procedure on Affymetrix microarray data.

### Laser Capture Microdissection

Breast tumor tissue was sectioned at 6 µm in a Leica CM 1850 Cryostat (Leica Microsystems GmbH). The sections were mounted on uncharged glass slides without the use of embedding media and placed immediately in 70% ethanol for 30 seconds. Subsequent dehydration was achieved using graded alcohols and xylene treatments as follows: 95 % ethanol for 1 minute, 100% ethanol for 1 minute (times 2), xylene for 2 minutes and second xylene 3 minutes. Slides were then dried in a laminar flow hood for 5 minutes prior to microdissection. Then, sections were laser captured microdissected with PixCell II LCM system (Arcturus Engineering). Approximately 5000 shots using the 30 micron infrared laser beam will be utilized to obtain approximately 10,000 cells per dissection. All samples were microdissected in duplicate on sequential sections.

### SDS-PAGE and In-gel Digestion

All membranes containing the microdissected cells from breast tumor tissue were removed and placed directly into a 1.5 mL Eppendorf tube. Membranes containing the microdissected cells were suspended in 20 µL of SDS sample buffer, reduced with DTT and heated in a 70-80°C water bath for approximately 10 min. The supernatant was then electrophoresed approximately 2 cm into a 10% Bis Tris gel, stained with Colloidal Blue with destaining with water, and the region was excised and subjected to in-gel trypsin digestion using a standard protocol. Briefly, the gel regions were excised and washed with 100 mM ammonium bicarbonate for 15 minutes. The liquid was discarded and replaced with fresh 100 mM ammonium bicarbonate and the proteins reduced with 5 mM DTT for 20 minutes at 55° C. After cooling to room temperature, iodoacetamide was added to 10 mM final concentration and placed in the dark for 20 minutes at room temperature. The solution was discarded and the gel pieces washed with 50% acetonitrile/50 mM ammonium bicarbonate for 20 minutes, followed by dehydration with 100% acetonitrile. The liquid was removed and the gel pieces were completely dried, reswelled with 0.5 µg of modified trypsin (Promega) in 100 mM NH_4_HCO_3,_ and digested overnight at 37°C. Peptides were extracted by three changes of 60% acetonitrile/0.1% TFA, and all extracts were combined and dried *in vacuo*. Samples were reconstituted in 35 µL 0.1 % formic acid for LC-MS/MS analysis.

### LC-MS/MS Analysis, Protein Identification and Quantitation

Peptide digests were analyzed on a Thermo LTQ Orbitrap Velos ion trap mass spectrometer equipped with an Eksigent NanoLC 2D pump and AS-1 autosampler as described previously [47]. Peptide sequence identification from MS/MS spectra employed the RefSeq Human protein sequence database, release version 54, and both database and peptide library search strategies [47]. For initial protein assembly, peptide identification stringency was set at a maximum of 1% reversed peptide matches, *i*.*e*., 2% peptide-to-spectrum matches (PSM) FDR and a minimum of 2 unique peptides to identify a given protein within the full data set. To minimize false-positive protein identifications, only proteins with a minimum of 6 matched spectra were considered. The full dataset contained 850,847 filtered spectra corresponding to 31,594 distinct spectrum-peptide sequence matches, which mapped to 24,946 distinct peptide sequences and 2,230 indistinguishable protein identifications. The protein-level FDR for the final assembly was 5.14%. Spectral counts for each protein in the final assembly were calculated as the sum of peptide-spectrum matches that met the criteria described above.

### Orthotopic xenograft studies

The human samples used to generate patient-derived xenograft (PDX) tumors, as well as the human non-tumor samples, were previously described [25]. The generation of the MTBTOM tumor model has been previously described [26]. 4-week-old WT FVB/N and immunocompromised NOD/SCID-gamma (NSG) female mice were purchased from Taconic Biosciences. Viably frozen MTB-TOM, HCI002, HCI009 and HCI010 tumor samples were transplanted into the mammary fat pad, following clearance of associated lymph node and epithelium, of respective FVB/N and NSG mice. Tumor growth was monitored daily by caliper measurement in two dimensions. When tumors reached 1 cm (MTBTOM) or 2 cm (PDX) in any dimension mice were euthanized, tumor and NAT were isolated, and flash-frozen in liquid nitrogen. The protocols described in this and other sections regarding animal studies were approved by the UCSF Institutional Animal Care and Use Committee. For the HCC1143 and HS578T control and Cx31 partial expression loss orthotopic xenografts, and for the BT549 shRNA knockdown orthotopic xenografts, 5 × 10_5_ cells were resuspended 1:1 with matrigel (Corning) and injected into the cleared mammary fat pads of 4-week-old WT NSG female mice. Tumor incidence and growth were monitored daily via palpation and caliper measurement, respectively. Mice were euthanized after 180 days or after tumors reached 2cm in any dimension. For HCC1143 *GJB3*^*WT*^ and *GJB3*^*Med*^ xenografts, a central slice of tumor and surrounding NAT was fixed in 4% paraformaldehyde and embedded in paraffin for histological sectioning, H&E staining and adipocyte area quantification, while the remaining tumor and NAT tissues were flash-frozen in liquid nitrogen. For other xenografts, NAT was isolated and flash-frozen in liquid nitrogen. For the CL316243 experiment, mice were randomized into experimental groups immediately post-orthotopic transplant. The following day, drug treatment was initiated and mice received vehicle or 1 mg/kg CL316243, delivered by intraperitoneal injection, daily until tumor incidence was recorded via palpation. For the Cx31 shRNA knockdown experiments, mice were randomized into experimental groups immediately post-orthotopic xenograft, and mice in the shCx31 knockdown group were administered doxycycline dietarily.

### Immunoblot analysis

Proteins were extracted using RIPA buffer (Thermo) and proteinase (Roche) plus phosphatase (Roche) inhibitor cocktails. Protein extracts were resolved using 4–12% SDS-PAGE gels (Life Technologies) and transferred to nitrocellulose membranes (Life Technologies). Membranes were probed with primary antibodies overnight on a 4 °C shaker, then incubated with horseradish peroxidase (HRP)-conjugated secondary antibodies, and signals were visualized with ECL (Bio-Rad). The primary antibodies targeting the following proteins were used: β-actin (actin) (sc-47778 HRP, Santa Cruz, 1:10,000), pHSL S563 (4139, Cell Signaling, 1:1000), HSL (4107, Cell Signaling, 1:1000), HNF4α (ab41898, Abcam, 1:1000), and Cx31 (ab156582, Abcam, 1:1000). Chemiluminescent signals were acquired with the Bio-Rad ChemiDoc XRS+ System equipped with a supersensitive CCD camera. Where indicated, unsaturated band intensities were quantified using Bio-Rad Image Lab software.

### Cell culture and virus production

A panel of established TN and RP human breast cancer cell lines, and their culture conditions, have previously been described [48]. No cell line used in this paper is listed in the database of commonly misidentified cell lines that is maintained by the International Cell Line Authentication Committee (ICLAC) (http://iclac.org/databases/cross-contaminations/). All lines were found to be negative for mycoplasma contamination. Lentiviruses for Cas9 and sgRNAs were produced in 293T cells using standard polyethylenimine (Polysciences Inc.) transfection protocols.

### Dye transfer and FACS analysis

For cancer cell-cancer cell transfer, monolayers of indicated lines (donors) were labelled with 1µM CalceinAM dye (Life Technologies) at 37°C for 40 min. Dye-loaded ‘donor’ cells were washed three times with PBS, and then single-cell suspensions of 1.5 × 10^5^ mCherry-labelled cells (recipients) were added for 5 hours. For CBX treatment studies, monolayers of indicated lines (recipients) were pre-treated for 24 hours with 150uM CBX or vehicle. Indicated ‘donor’ cells were loaded in suspension with CalceinAM dye (Life Technologies) at 37°C for 40min, washed three times with PBS, and added onto indicated ‘recipient’ cells for 5 hours. Dye transfer was quantified by BD LSRFORTESSA or BD LSR II (BD Biosciences). Gating strategy to identify mCherry-positive, Calcein-positive cell population is described in Figure S4. For cancer cell-adipocyte transfer, monolayers of indicated control or Cx31 partial knockout lines (donors) were labelled with 1 µM CalceinAM dye at 37°C for 40 min. Dye-loaded cells were washed three times with PBS, and then primary mammary adipose tissues (recipient) were added for 5 hours. Primary adipose tissue was isolated from co-culture, washed with PBS, and dye transfer was quantified by measurement of total adipose fluorescence using a Tecan fluorescent plate reader.

### Gene expression analysis

TCGA breast-invasive carcinoma data set was sourced from data generated by TCGA Research Network (http://cancergenome.nih.gov), made available on the University of California, Santa Cruz (UCSC) Cancer Browser. Single-cell RNAseq data was sourced from data generated by Chung, et al.[34]. For the MTBTOM data set, 11 endpoint MTBTOM orthotopic xenografts generated as described above, and 3 mammary glands from naïve mice, were flash-frozen in liquid nitrogen. Library preparation and Illumina RNAseq was performed by Q^2^Solutions (www.q2labsolutions.com). Gene expression analyses were performed using the ‘limma’ R package [49]. For the panel of established TN and RP human breast cancer cell lines [48], library preparation and Illumina RNAseq was performed by Novogene (www.novogene.com). All RNA was isolated using the RNAeasy kit (Qiagen).

### ATP quantification

To determine the effects of CBX treatment on ATP levels, tumor cells were seeded in 96-well plates at 5,000–7,000 cells per well and cultured in the presence of 0 or 150 μM CBX (Sigma) for 24 hours, with triplicate samples for each condition. Relative ATP concentrations were determined using the CellTiter-Glo Luminescent Cell Viability Assay (Promega).

### Isolation of primary mammary adipose tissue

Anonymous reduction mammoplasty samples were acquired from the Cooperative Human Tissue Network (CHTN). Samples were washed in DPBS supplemented with 1 % Penicillin/Streptomycin and 0.1 % Gentamicin (all GIBCO). Mammary adipose tissue was separated mechanically from epithelial tissue using a razor blade, and was then cryopreserved in freezing medium (10% DMSO (Sigma) in FBS (X&Y Cell Culture)).

### Immunofluorescence staining and microscopy

For adipose tissue cancer cell co-cultures imaged whole mount, 2 × 10^6^ of the indicated GFP-labelled cell line was suspended in 500uL DMEM/F-12(Gibco 11320033) containing 10% FBS and injected into primary mammary adipose tissue from a healthy individual, then cultured at 37°C for 24 hours. For immunofluorescence labeling of co-culture tissues, samples were washed three times in PBS and fixed in 4% paraformaldehyde, permeabilized in 0.5% Triton X-100 for 15min, and blocked in 10% goat serum in PBS with 0.25g/L BSA, 0.2% Triton X-100, and 0.41% Tween-20 overnight. Samples were then incubated overnight with primary antibodies (Cx31, WH0002707M1, Sigma, 1:100, and pHSL(S563), 4139, Cell Sig, 1:100), and then overnight with Alexa Fluor-647 or -546 conjugated antibodies. Finally, using an established protocol for whole mount breast tissue imaging[37], co-culture tissues were cleared through overnight incubation at 4 °C in a ‘FUnGI’ solution of 50% glycerol (vol/vol), 2.5 M fructose, 2.5 M urea, 10.6 mM Tris Base, and 1 mM EDTA. Confocal images were acquired using a Zeiss LSM900 with Airyscan 2 detector. For pHSL(S563) image quantification, fluorescence was measured using Fiji imaging software, and Difference of Gaussians was used for analysis of puncta number and percent area. For sectioned adipose tissue coculture, 1 × 10^6^ of the indicated mCherry-labelled cell line was injected into primary mammary adipose tissue and cultured at 37°C for 18 hours. The co-cultures were examined using fluorescent microscopy to identify regions of adipose tissue containing mCherry-positive cancer cells. These regions were isolated and fixed in 4% paraformaldehyde and embedded in paraffin. Primary TNBCs used for immunofluorescence were identified and retrieved from the clinical archives of the University of California San Francisco (UCSF) Department of Pathology. All tumors consisted of estrogen receptor (ER)-, progesterone receptor (PR)-, and HER2-negative invasive ductal carcinomas. Breast tissue was fixed in 10% formalin and embedded in paraffin. Tumor blocks with sufficient tumor and adjacent (at least 0.5 cm) normal tissue were selected, and 4μm sections were cut on plus-charged slides for immunofluorescence. This study was approved by the UCSF institutional review board. For immunofluorescence labeling of sectioned co-cultures and primary TNBC, slides were dewaxed in xylene followed by rehydration in graded ethanol (100, 95, 70%) and deionized H2O. Antigen retrieval was performed in 10mM Tris, 1mM EDTA, 0.05% Tween 20, pH 9 at 121 °C for 4 min. Subsequently, tissue sections were blocked in 1% bovine serum albumin, 2% fetal bovine serum in PBS for 5 min, and incubated with primary antibodies (Cx31, 12880, Proteintech, 1:50 and pan-cytokeratin, sc-81714, Santa Cruz, 1:50) overnight at 4 °C. Following several PBS washes, sections were incubated with Alexa Fluor-488 or - 568 conjugated antibodies, counterstained with DAPI (Sigma), and mounted using Vectashield (Vector). Epifluorescence images were acquired either by spinning disk microscopy on a customized microscope setup as previously described [50-52] except that the system was upgraded with a next generation scientific CCD camera (cMyo, 293 Photometrics) with 4.5 μm pixels allowing optimal spatial sampling using a Å∼60 NA 1.49 objective (CFI 294 APO TIRF; Nikon), or at the UCSF Nikon Imaging Center using a Nikon Ti Microscope equipped with an Andor Zyla 5.5 megapixel sCMOS camera and Lumencor Spectra-X 6-channel LED illuminator. Images were collected using a Plan Apo λ 20x / 0.75 lens.

### Generation of Cx31 partial expression loss lines

LentiCas9-Blast (Addgene plasmid #52962) and lentiGuide-Puro (Addgene plasmid #52963) were gifts from Feng Zhang. sgRNAs against Cx31 were constructed using the Feng Zhang Lab CRISPR Design Tool (crispr.mit.edu). sgRNAs used were as follows:

Cx31 exon 1 sg1: CCAGATGCGCCCGAACGCTGTGG (HS578T *GJB3*^*Med-1*^ and HCC1143 *GJB3*^*Med*^)

Cx31 exon 1 sg2: CCGGGTGCTGGTATACGTGGTGG (HS578T *GJB3*^*Med-2*^ and HCC1143 *GJB3*^*Low*^)

ShRNAs against Cx31 and GFP control were constructed using Tet-pLKO-Puro (Addgene plasmid #21915). shRNAs used were as follows:

shCx31:

Cx31.shRNA3_forward: ccggAAGCTCATCATTGAGTTCCT CctcgagGAGGAACTCAATGATGAGCTTtttttg Cx31.shRNA3_reverse: aattcaaaaaAAGCTCATCATTGAGTTCCTCctcgagGAGGAACTCAATGATGAGCTT

shGFP [53]:

shGFP_forward: CCGGTACAACAGCCACAACGTCTATCTCGACATAGACTTGTGGCTGTTGTATTTTTG shGFP_reverse: CAAAAATACAACAGCCACAACGTCTATGTCGAGATAGACGTTGTGGCTGTTGTACCGG

Lentiviral transduction was performed in DMEM supplemented with 10% FBS and polybrene 10 μg/mL. For sgRNA transduction, Cas9-expressing cells were enriched by Blasticidin (10-15 μg/mL Gemini BioProducts) selection for seven days. Cas9+ cells were subsequently transduced with lentiGuide-Puro (with sgRNAs targeting Cx31) followed by puromycin (1 μg/mL; Gibco) for seven days. Thereafter, clonal selection was performed and clones screened for loss of target gene protein expression by immunoblot analysis. For shRNAs, cells were transduced with Tet-pLKO-Puro (with shRNAs targeting Cx31 or GFP control[53]) followed by puromycin (2 ug/mL; Gibco) for seven days, after which knockdown of target protein was confirmed by immunoblot analysis.

### cAMP quantification

For *in vitro* studies, tumor cells were seeded in 96-well plates at 5,000–7,000 cells per well and cultured in the presence of 0 or 150 μM CBX (Sigma) for 24 hours, with triplicate samples for each condition. Changes in cAMP concentration were determined using the cAMP-Glo Assay (Promega).

For *in vivo* studies, frozen tissue was homogenized using a TissueLyser in 300 μl of 40:40:20 acetonitrile:methanol:water with the addition of 1 nM (final concentration) of D3-[15N]serine as an internal extraction standard (Cambridge Isotopes Laboratories Inc, DNLM-6863). 10 μl of cleared supernatant (via centrifugation at 15,000 r.p.m., 10 min, at 4 °C) was used for SRM–LC-MS/MS using a normal-phase Luna NH2 column (Phenomenex). Mobile phases were buffer A (composed of 100% acetonitrile) and buffer B (composed of 95:5 water:acetonitrile). Solvent modifiers were 0.2% ammonium hydroxide with 50 mM ammonium acetate for negative ionization mode. cAMP levels were analyzed using the MassHunter software package (Agilent Technologies) by quantifying the transition from parent precursor mass to product ions.

### cAMP transfer

For cancer cell-adipocyte transfer, monolayers of indicated control or Cx31partial knockout lines (donors) were labelled with 2µM fluo-cAMP (Biolog Life Science Institute) at 37°C for 30 min. cAMP-loaded cells were washed three times with PBS, and then primary mammary adipose tissues (recipient) were added for 5 hours. Primary adipose tissue was isolated from co-culture, washed with PBS, and cAMP transfer was quantified by measurement of total adipose fluorescence using a Tecan fluorescent plate reader.

### Preadipocyte differentiation and qRT-PCR

Primary mouse preadipocytes were differentiated as previously described [54]. Monolayers of differentiated adipocytes were washed with PBS, and then treated with vehicle or 10μM forskolin (Sigma), or seeded with 1 × 10^5^ of the indicated cancer lines. Total RNA was isolated from co-cultures after 20 hours using the RNeasy kit (Qiagen). One μg of total RNA was reverse transcribed using iScript cDNA synthesis kit (Bio-Rad). The relative expression of UCP1, aP2, and GAPDH was analyzed using a SYBR Green Real-Time PCR kit (Thermo) with an Applied Biosystems QuantStudio 6 Flex Real-Time PCR System thermocycler (Thermo). Variation was determined using the ΔΔCT method [55](*48*) with GAPDH mRNA levels as an internal control. Mouse-specific primers used were as follows:

GAPDH forward: CCAGCTACTCGCGGCTTTA

GAPDH reverse: GTTCACACCGACCTTCACCA UCP1

forward: CACCTTCCCGCTGGACACT

UCP1 reverse: CCCTAGGACACCTTTATACCTAATGG

aP2 forward: ACACCGAGATTTCCTTCAAACTG

aP2 reverse: CCATCTAGGGTTATGATGCTCTTCA

### Proliferation assays

To determine the effects of Cx31 partial knockout on cell proliferation and viability, the indicated cell lines were seeded in 6-well plates at 1.5 × 10^5^ cells/well. Cells were harvested at 24, 48 and 72 h. Cell counts and cell viability by trypan blue exclusion were determined using the Countess Automated Cell Counter (Life Technologies) according to the manufacturer’s instructions.

### Statistical analysis

Prism software was used to generate and analyze Spearman correlation (Fig. 1D) and the survival plots (Figs. 4B, 4C and 4G). Correlation *P* values were generated using ordinary one-way ANOVA (Figs. 1H, 3F, 3H, 4D and 4F), ordinary one-way ANOVA with multiple comparisons (Fig. 2B), repeated measures one-way ANOVA (Figs. 1B, 1F, 3E, and 3G), repeated measures mixed effects model (Figs. 1C, 1E and 1G), and unpaired two-tailed t test (Figs. 4A, 4E, S1A, S1B). Survival plot *P* values was generated using a log-rank test. All differential expression analyses (Fig. 2D and 2F) were done using the ‘limma’ R package [49].

**Fig. 1.**
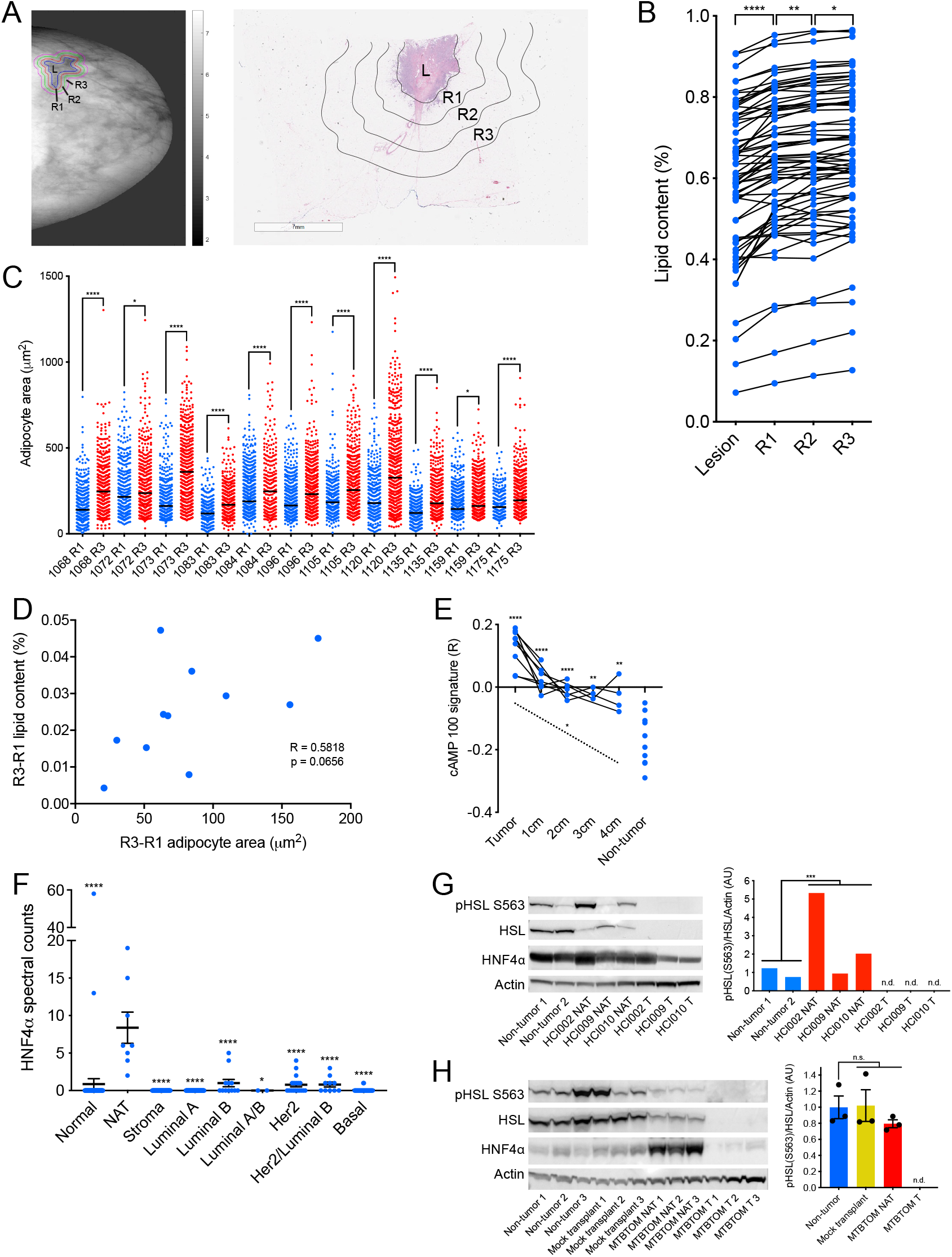
Lipolysis and lipolytic signaling are activated in breast tumor-adjacent adipocytes from breast cancer patients and mouse models of breast cancer. (A) Representative lipid content image (left) and hematoxylin and eosin stained excision specimen (right) from patients with invasive breast cancer. The lesion (L), and NAT 0-2 mm (R1), 2-4 mm (R2), and 4-6 mm (R3) away are indicated. (B) Percent lipid content (lipid content / lipid + water + protein content) of L, R1, R2 and R3 from patients (n = 46) with invasive breast cancer. (C) Adipocyte area in R1 and R3 from a subset of patients (n = 11) in B. The black line indicates mean adipocyte area, and each patient identifier is indicated. Each point represents individual adipocyte. (D) Correlation of change in lipid content in B and change in average adipocyte area in C from R3 to R1 for matched patients in C. Spearman correlation and two-tailed t test were used to generate the correlation coefficient and associated P value. (E) ssGSEA enrichment scores for cAMP-dependent lipolysis signature in primary breast tumors (n = 9), NAT 1 cm (n = 7), 2 cm (n = 5), 3 cm (n = 3), and 4 cm (n = 4), and healthy non-tumor breast tissue (n = 10). Dotted line indicates fixed effects analysis across matched samples. (F) HNF4a peptide counts from LC-MS/MS of primary tissue from healthy control breast tissue (n = 42), NAT (n = 4), stroma (n = 36), and luminal A (n = 38), luminal B (n = 6), luminal A/B (n = 1), HER2-amplified (n = 9), HER2-amplified/luminal B (n = 5), and basal (n = 16) tumors. Each point represents individual sample LCM on which LC-MS/MS was performed. LCM and LC-MS/MS was performed in technical duplicate on sequential histological slides from each patient. (G) Immunoblot analysis (left) showing expression levels of lipolysis activators HSL and HNF4a, and phosphorylated HSL (pHSL S563) in healthy non-tumor mammary gland and NAT and tumor tissues from a panel of PDXs. Quantification (right) of pHSL/HSL ratio, normalized to b-actin levels. (H) Immunoblot analysis (left) showing expression levels of lipolysis activators HSL and HNF4a, and phosphorylated HSL (pHSL S563) in healthy non-tumor mammary gland, mock-transplanted mammary gland, and NAT and tumor tissues from MTBTOM allografts. Quantification (right) of pHSL/HSL ratio, normalized to b-actin levels. For (B) and (E) black lines indicate matched samples from individual patients. For (F) and (H) mean ± s.e.m. is shown. *P < 0.05, **P < 0.01, ***P < 0.001, ****P < 0.0001; repeated measures one-way ANOVA (B) and (F), repeated measures mixed effects model (C), (E), and (G), ordinary one-way ANOVA (H).

**Fig. 2.**
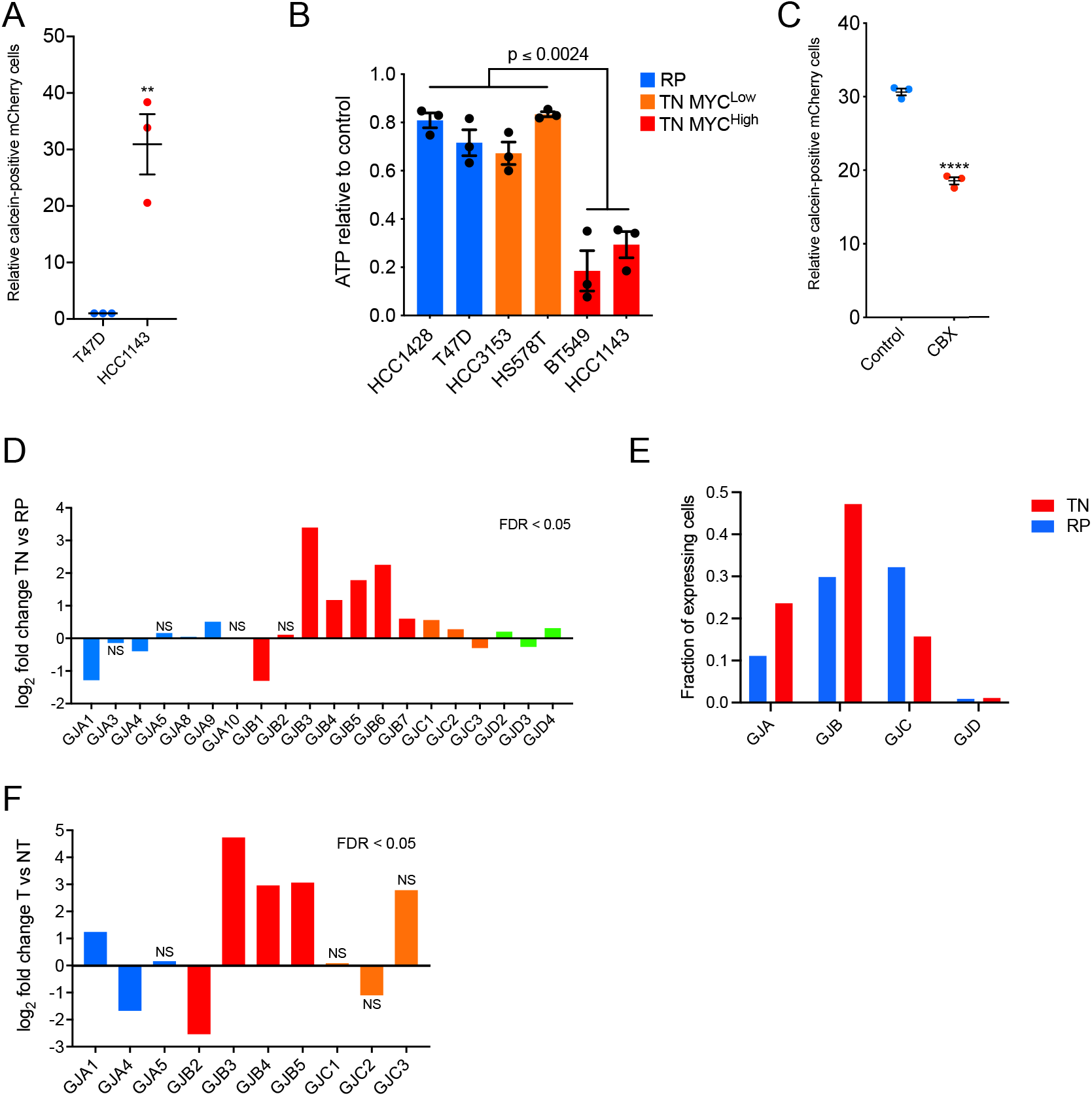
Breast cancer cells form functional gap junctions and express Cx31. (A) Relative frequency of dye transfer from Calcein AM-loaded cells (donor) to mCherry-labelled cells (recipient) as determined by FACS (fluorescence-activated cell sorting) analysis. (B) ATP levels in TN high MYC, TN low MYC, and RP cell lines after treatment with 150 μM CBX for 24 hours relative to untreated (control) cells. (C) Relative frequency of dye transfer from Calcein AM-loaded cells (donor) to mCherry-labeled cells (recipient) treated with 150uM CBX or vehicle control for 24 hours, as determined by FACS analysis. (D) Fold change (log2) in expression of indicated connexin genes in TN (n = 123) versus RP (n = 648) tumors based on RNA-seq data acquired from TCGA of 771 breast cancer patients. (E) Fraction of cells in (n=11) patient tumors of RP and TNBC subtypes expressing indicated gap junction (GJ) family members, based on sc-RNA-seq of 317 tumor cells. (F) Fold change (log2) in expression of indicated connexin genes in T (n = 11) versus NT (n = 3) tissues based on RNA-seq data from MTBTOM allograft-bearing mice or healthy controls, respectively. For (A) and (C) mean ± s.e.m. of three independent biological replicates is shown. **P < 0.01, ****P < 0.0001; unpaired two-tailed t test (A) and (C), ordinary one-way ANOVA (B). For (D) and (F), all differential expression analysis was done using the ‘limma’ R package.

## Code availability

Publicly available data sets were acquired as noted. Our annotations of the TCGA data set are available (https://bitbucket.org/jeevb/brca).

## Main text

A variety of cancers, including those of the breast, arise near or within adipose tissue depots [1]. Therefore, during tumor development a heterotypic cell-cell interface exists between adipocytes and cancer cells in these organs. We and others have demonstrated that triple-negative breast cancers (TNBC, estrogen/progesterone/HER2 receptor-negative) utilize and require fatty acid oxidation to fuel bioenergetic metabolism [2, 3]. The origin of fatty acids which meet this necessity remains largely unclear. Adipocyte lipolysis has been linked by several studies to secretion of pro-tumorigenic cytokines by cancer-associated adipocytes, including elevation of pro-inflammatory signals such as tumor necrosis factor-α [4-11]. Multiple models provide evidence that these adipocyte-derived fatty acids can be taken up and oxidized by proximate cancer cells [4-10, 12]. These studies, however, have widely modeled the cancer-adipocyte interface *in vitro* using transwell co-culture methods that cannot recapitulate the direct cell-cell contact observed *in vivo* [6-9, 11-13]. Furthermore, clinical evidence for elevated lipolysis in breast tumor-adjacent adipocytes has not been well established. Mammary adipocytes undergo enhanced lipolysis when in close proximity to non-tumor epithelial cells, suggesting that local prolipolytic mechanisms exist, but have yet to be identified between tumor cells and adipocytes [5, 14]. Thus, we set out to study the breast cancer-adipocyte interface and determine the contribution of cell-cell contact to tumorigenesis.

To determine if lipolysis occurs in normal tissue adjacent to breast tumors (NAT) which includes adipocytes, we employed four independent strategies. First, we employed three-component breast (3CB) composition measurement, a radiographic imaging method derived from dual-energy mammography that allows for quantification of a tissue’s water, lipid and protein content [15]. We postulated that, if tumors induce lipolysis in adipocytes, we will observe differences in lipid content between NAT nearer the tumor and NAT farther away. Using 3CB imaging, we assessed the lipid content of breast tumors and the first 6 mm of surrounding NAT, segmented into 2 mm “rings,” from 46 patients with invasive breast cancer (Fig. 1A and Table S1). As we have previously demonstrated [16], we found a significant decrease in lipid content in tumor lesions compared to NAT 0-2 mm away (R1) (Fig. 1B). This difference is congruent with breast tumors being epithelial in nature, while adipose tissue is the major constituent of normal breast [14]. Remarkably, we also found that within NAT there was a significant stepwise decrease in lipid content comparing R3 (4-6 mm) to R2 (2-4 mm), and R2 to R1 (Fig. 1B). In addition, we asked whether changes in lipid content between R3 and R1 NAT correlate with receptor status or tumor grade (Table S1). We found that NAT surrounding triple-negative (TN) and grade 2/3 tumors trended towards a greater average decrease in lipid content between R3 and R1 than NAT surrounding receptor-positive (RP) and grade 1 tumors, respectively (Fig. S1, A and B). These data suggest that adipocytes near breast tumors have partially depleted lipid stores, and that TN and higher-grade tumors may induce this phenomenon to a greater degree than RP and low-grade tumors. We quantified average adipocyte size in R1 and R3 in the 11 of the 46 patients imaged with 3CB for whom we had access to histological sections of treatment-naïve tumor and NAT at the time of surgical resection (Fig. 1A, Fig. S1C and Table S1). Similar to the change in lipid content observed with 3CB, we found a significant decrease in adipocyte size in R1 compared to R3 in all patients analyzed, suggesting adipocytes are smaller when nearer to breast tumors (Fig. 1C). Finally, we correlated the change in lipid content and adipocyte size on an individual patient basis. We found a marked positive correlation (R = 0.5818, p = 0.0656) between the change in lipid content and adipocyte area (Fig. 1D). Taken together, these data suggest adipocytes are smaller and have diminished lipid content, two phenotypes that are established indicators of lipolysis [17], when adjacent to breast tumors.

Second, we sought to determine if gene expression changes associated with lipolysis were observed in tumor-adjacent adipocytes. We generated a lipolysis gene expression signature by identifying the 100 genes most upregulated when a differentiated adipocyte cell culture model is stimulated with cAMP, a critical pro-lipolytic signaling molecule [18]. We then used a publicly available gene expression dataset for primary breast tumors as well as matched NAT 1, 2, 3 and 4 cm away, to determine if enrichment of the lipolysis signature occurred in NAT in comparison to non-tumor breast tissue obtained from healthy individuals using single-set gene set enrichment analysis [19, 20]. We found a significant elevation of the cAMP-dependent lipolysis signature in tumor and NAT from all analyzed regions compared to control tissue (Fig. 1E). These data indicate that lipolytic signaling is activated in breast-tumor adjacent adipocytes up to 4 cm away from the primary tumor. While adipose tissue is sparsely innervated, a recent study found that adipocytes can propagate pro-lipolytic sympathetic signals via direct transfer of cAMP through adipocyte-adipocyte gap junctions [21]. We observed elevation of cAMP signaling up to 4 cm away from patient tumors (Fig. 1E), suggesting that tumor-adjacent adipocytes might also disperse a pro-lipolytic stimulus to distant adipocytes via gap junctions.

Third, we sought to determine if there are changes to protein abundance in tumor-adjacent NAT indicative of lipolysis activation. We conducted laser capture microdissection (LCM, 10,000 cells per capture) on primary breast tumors from 75 patients, representing all major PAM50 subtypes. For a subset of patients, we also collected matched stroma and/or NAT. As a control, we conducted LCM on non-tumor breast tissue from 42 healthy subjects (Table S2A). Global proteomic analysis was performed using liquid chromatography-tandem mass spectrometry (LC-MS/MS) (Table S2B). Notably, one of the most significantly upregulated proteins in NAT, and indeed one of the most NAT-specific proteins, compared to all other tissues examined was hepatocyte nuclear factor 4-α (HNF4α) (Fig. 1F). As HNF4α is an established, essential activator of lipolysis in adipose tissue [22], these data indicate lipolysis is robustly activated in breast-tumor adjacent adipose tissue.

Fourth, we sought to validate the observations made in our clinical datasets using mouse models of breast cancer. Hormone sensitive lipase (HSL) is a critical lipolytic enzyme; its activation by cAMP-dependent protein kinase A (PKA) leads to phosphorylation at serine 563 [17, 18], while prolonged activation results in down-regulation of total HSL expression through a negative feedback mechanism [23, 24]. We performed immunoblot analysis to probe for HSL, phospho-HSL (S563) and HNF4α in tumor and NAT, as well as corresponding control mammary tissues, from three well-characterized breast cancer patient-derived xenograft (PDX) models (HCI002, HCI009, HCI010) and a transgenic model of MYC-driven TNBC (MTB-TOM) [25, 26]. In all models analyzed, a downregulation of total HSL in NAT compared to control tissue was observed (Fig. 1, G and H). Downregulation of total HSL has been observed in obesity and in an independent analysis of primary breast tumor NAT, and is thought to be the result of a negative feedback loop in adipocytes in response to chronic lipolysis [23, 24]. Additionally, in 3 of the 4 models examined we found an increase in HNF4α protein or in phospho-HSL/total HSL ratio (Fig. 1, G and H), both characteristic of increased lipolysis [17, 22]. Taken together, our concurrent findings in 3 independent clinical datasets and several models of patient-derived and transgenic breast cancers in mice indicate that lipolysis is activated, to varying degrees, in breast cancer-adjacent adipose tissue. These findings support the conclusion that “normal” tissue adjacent to tumors is, in fact, not normal [27]; in the context of breast cancer, tumor-adjacent adipocytes have markers of activated lipolysis with corresponding diminished lipid stores.

We next sought to determine the contribution of cell-cell contact to lipolysis activation in breast tumor-adjacent adipocytes. Gap junctions are cell-cell junctions formed by a family of proteins called connexins, which are known to transport a variety of small molecules (<1 kD), including cAMP [21, 28]. Connexins were long thought to play tumor-suppressive roles in cancer, but recent evidence from a variety of tumor types has challenged this notion [28-31]. Given that adipocytes are capable of transferring cAMP and activating lipolysis in a homotypic interaction with other adipocytes[21], we hypothesized that gap junctions may also form between tumor cells and adipocytes in a heterotypic fashion to activate lipolysis via transfer of cAMP. Using a well-established dye transfer assay [30], we first probed for functional gap junction formation between breast cancer cells. We tested whether the TNBC cell line HCC1143 or the more indolent RP cell line T47D could transfer gap-junction dependent dyes to the same tumor cell line. Both lines formed functional gap junctions, but dye transfer between HCC1143 cells was 30-fold increased (Fig. 2A) compared to transfer amongst T47D cells. Thus, we reasoned there may be differences in sensitivity to gap junction inhibition between TN and RP cells. Furthermore, given the upregulation of MYC in the majority of TNBC [32, 33], we asked whether MYC expression affects gap junction dependence. We examined if gap junction inhibition alters cellular ATP as a proxy for cell abundance in a panel of TN and RP human breast cell lines with varying MYC levels [2]. Intriguingly, TNBC cell lines with high MYC expression [2], including HCC1143, were significantly more sensitive to 24 hours of treatment with the pan-gap junction inhibitor carbenoxolone (CBX) than the low MYC TNBC or RP cell lines tested (Fig. 2B). In addition, dye uptake in HCC1143 cells was significantly reduced (30.63%, p<0.0001) following treatment with CBX (Fig. 2C). These data suggest that gap junction communication occurs between breast cancer cells, and that a threshold amount of gap junction activity may be required for high MYC TN cell viability.

To delineate the role of connexins in TN compared to RP breast cancer further, we examined the expression of the 21 connexin genes in 771 primary human breast cancers, TN (*n* = 123) and RP (*n* = 648), using publicly available RNA-seq data from The Cancer Genome Atlas (TCGA). Of the 20 connexins for which data was available, 5/20 were significantly downregulated, and 11/20 were significantly upregulated. These 11 upregulated connexins included 5 of the 7 gap junction B (GJB) family members (Fig. 2D). To probe gap junction expression at the cellular level, we also examined scRNA-seq (n=317) of primary patient tumors (n=11) [34]. Expression of GJBs was observed in a greater fraction (47.2% vs. 29.8%) of TN than RP tumor cells, and GJBs were the most frequently expressed gap junction family for TN, but not for RP tumor cells (Fig. 2E and Fig. S2). As an independent approach to examine *in vivo* expression of connexins in TNBC, we then performed RNA-seq on MTBTOM tumors and non-tumor control tissue (Table S3). Of the 10 connexins for which data were available, 2/10 were significantly downregulated, 4/10 were significantly upregulated, and 4/10 were not significantly changed in MTBTOM tumors versus control tissue (Fig. 2F). Connexin 31 (*GJB3*, Cx31) was the most significantly elevated connexin in both human TN tumors and the MYC-driven TNBC model. Thus, we focused the remainder of our studies on Cx31. Cx31 has been found to be expressed in keratinocytes, the small intestine, and the colon [35, 36]. Although roles for various connexins as oncogenes and/or tumor suppressors have been described [28, 29], a pro-tumorigenic function of Cx31 has not been established.

Accordingly, we sought to determine if functional Cx31-containing gap junctions form between breast cancer cells and adipocytes. To validate the presence of cancer-adipocyte gap junctions in TNBC, we began by examining primary patient biopsies for expression of Cx31 and of pan-cytokeratin to distinguish epithelial tumor cells. We found that both TN tumor cells and adipocytes robustly express Cx31 at the plasma membrane. Further, we found many points of cell-cell contact occurred *in vivo* between tumors and adipocytes (Fig. 3A). To model the cell-cell contact observed *in vivo* between breast cancer cells and adipocytes, we developed three independent co-culture models. First, we performed 3-dimensional *ex vivo* studies by coculturing breast cancer cells directly within primary patient breast fat (Fig. 3B). We stably transduced HCC1143 (TNBC) and T47D (RP) with an mCherry expression plasmid, then injected either mCherry-HCC1143 or -T47D cells directly into mammary adipose tissue (WD43177) and co-cultured overnight. Tumor cell-adipocyte co-cultures were formalin-fixed, paraffin-embedded and probed for Cx31 and pan-cytokeratin expression, then imaged using immunofluorescent microscopy. We found that both HCC1143 cells and adipocytes robustly expressed Cx31 at the plasma membrane; HCC1143 formed close cell-cell contacts with primary adipocytes (Fig. 3B, top). In contrast, while T47D cells formed cancer cell-cancer cell contacts, we did not observe close cancer cell-adipocyte contacts (Fig. 3B, bottom). These data suggest that Cx31 can be expressed at both the tumor cell and adipocyte plasma membrane, and that breast cancer cells can form close cell-cell contacts with adipocytes.

**Fig. 3.**
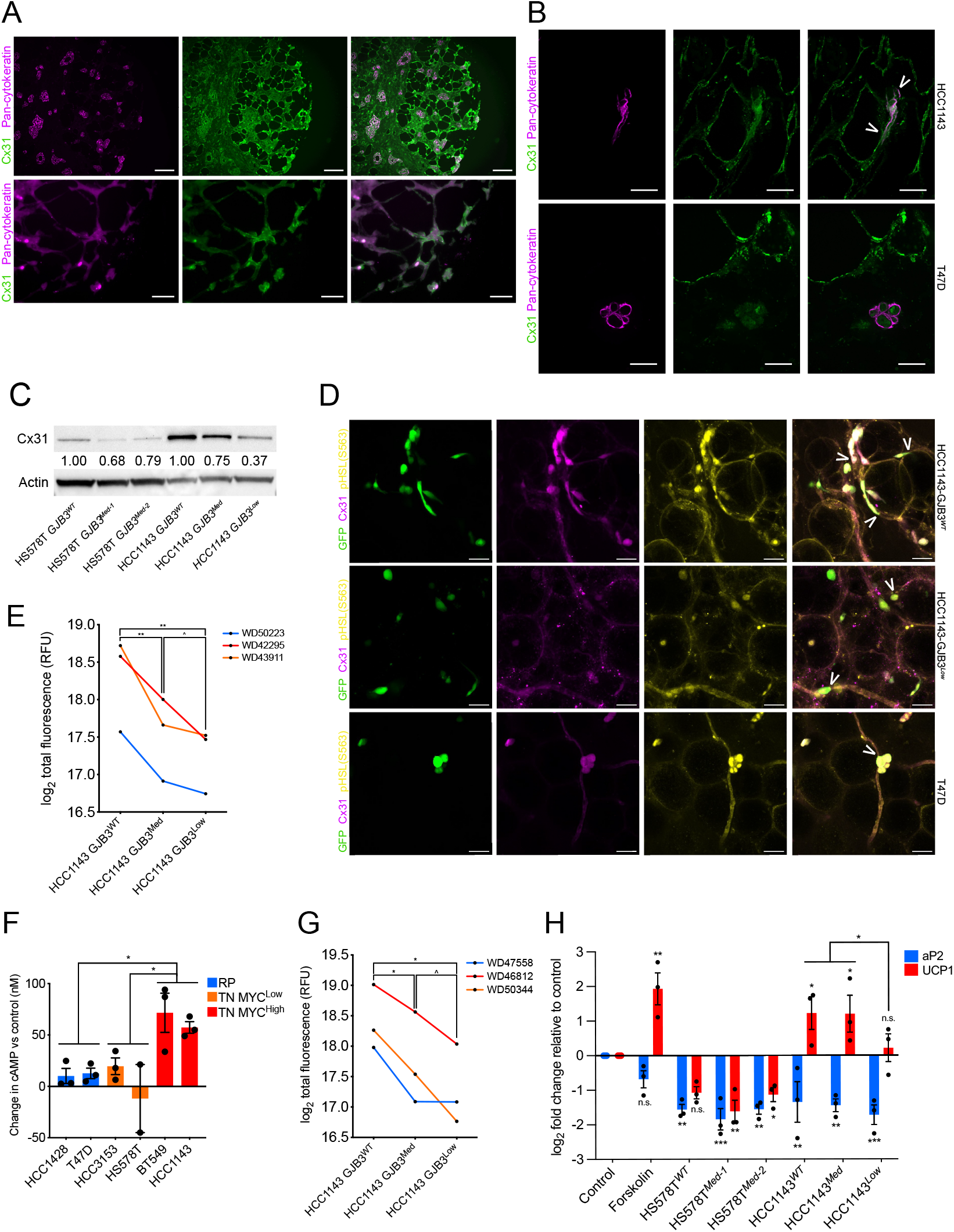
Breast cancer cell-adipocyte gap junctions form, transfer cAMP, and activate lipolytic signaling in a Cx31-dependent manner. (A) Staining with Cx31 (green) and pan-cytokeratin (magenta) of primary TNBC patient biopsies. Scale bar, top 100 μm, bottom 25 μm. (B) Staining with Cx31 (green) and pan-cytokeratin (magenta) of primary mammary tissue from a healthy individual (WD43177) injected with TN mCherry-HCC1143 cells and cocultured overnight. White arrowheads indicate co-staining of Cx31 with contact point between HCC1143 and adipocyte plasma membranes. Scale bar, 25 μm. (C) Immunoblot analysis showing protein expression levels of Cx31 in vitro in a panel of clonally derived control GJB3 WT and partial depletion TN lines with one-third and two-thirds loss of GJB3 expression. For the Cx31-depleted lines each clone is referred to by level of GJB3 expression (e.g. GJB3Med expresses two-thirds WT level, and GJB3Low expresses one third GJB3WT level). Quantification of Cx31 level normalized to b-actin level is indicated. (D) Staining with Cx31 (magenta) and pHSL(S563) (yellow) of primary mammary tissue from a healthy individual (WD49393) injected with GFP-expressing HCC1143-GJB3WT (top), HCC1143-GJB3Low (middle), or T47D cells (bottom) and co-cultured overnight. White arrowheads indicate co-staining of Cx31 and pHSL(S563) at contact point between GFP cancer cells and adipocytes. Scale bar, 50 μm. (E) Dye transfer from indicated HCC1143 control and Cx31-depleted lines to primary mammary adipose tissue of indicated (n = 3) healthy individuals. (F) cAMP levels in TN high MYC, TN low MYC, and RP cell lines after treatment with 150 μM CBX for 24 hours, relative to untreated (control) cells. (G) cAMP transfer from indicated HCC1143 control and Cx31 partial expression loss lines to primary mammary adipose tissue of indicated (n = 3) healthy individuals. (H) Fold change in UCP1 and aP2 expression in differentiated adipocytes after treatment with vehicle (control) or 10 μM forskolin, or co-cultured with indicated Cx31 partial expression loss lines for 24 hours. For (F) and (H) mean ± s.e.m. of three independent biological replicates is shown. ^P < 0.10, *P < 0.05, **P < 0.01, ***P < 0.001; repeated measures one-way ANOVA for (E) and (G), ordinary one-way ANOVA for (F) and (H).

To determine whether breast cancer cells rely upon Cx31-containing gap junctions to influence adipocyte function, we used CRISPR/Cas9 to generate a series of *GJB3* depleted TN lines (HS578T and HCC1143). In TN MYC-high TN cell line HCC1143, we generated two clones, with ∼1/3 and ∼2/3 *GJB3* expression loss (HCC1143 *GJB3*^*Med*^ and *GJB3*^*Low*^). In TN MYC-low line HS578T, we generated two distinct clones with ∼1/3 *GJB3* expression loss (HS578T *GJB3*^*Med-1*^ and *GJB3*^*Med-2*^). Despite several attempts we were unable to generate TN cell lines with complete Cx31 loss, strongly suggesting that a basal level of Cx31 expression is required for TN cancer cell growth.

To examine how Cx31 expression impacted cancer cell-adipocyte contact, we performed *ex vivo* co-cultures with primary patient breast fat using the partially depleted Cx31 cell lines. We stably transduced TN HCC1143 *GJB3*^*WT*^ and *GJB3*^*Low*^ cell lines, as well as RP line T47D with a GFP expression plasmid, then injected each line directly into primary mammary adipose tissue (WD49393). After overnight incubation, co-cultured tissues were formalin-fixed and probed for expression of Cx31 and lipolysis marker pHSL(S563) [17]. Tissues were then were cleared [37] and imaged via whole mount fluorescence microscopy. We found that HCC1143 *GJB3*^*WT*^ cells formed extended cancer cell-adipocyte contacts, in tight conformation with adjacent adipocytes (Fig. 3D, top). In contrast, Cx31-depleted HCC1143 *GJB3*^*Low*^ cells formed tangential contacts with adjacent adipocytes (Fig. 3D, middle), which we note closely mimic the tangential cancer cell-adipocyte conformation observed in T47D (RP) cocultures (Fig. 3D, bottom). In mock co-cultures, our positive control forskolin, which raises intracellular cAMP levels by activating adenylyl cyclase [18], robustly induced pHSL(S563) expression and increased puncta compared to vehicle-treated mammary adipose tissue (Fig. S3, A-C). We observed greater pHSL(S563) expression and elevated puncta in adipose tissue cocultured with HCC1143 *GJB3*^*WT*^ cells than tissues with HCC1143 *GJB3*^*Low*^ or T47D cells (Fig. S3, D and E), indicating more cAMP-dependent PKA activity. These results suggest that Cx31 level in breast cancer can moderate cell contact with surrounding adipocytes and alter lipolytic signaling.

We next sought to determine if Cx31 expression impacted tumor cell-adipocyte communication using a co-culture model in which HCC1143 *GJB3*^*WT*^, *GJB3*^*Med*^ or *GJB3*^*Low*^ cells were seeded in 2D culture and loaded with gap junction-transferable dye. We added primary mammary adipose tissue from three healthy individuals (WD42295, WD43911, WD50223) directly on top of the monolayers to permit direct contact. Tumor cells and adipocytes were co-cultured for 5 hours and then assayed for gap junction-dependent dye transfer from the cancer cells to adipocytes. We found that robust dye transfer occurred from the HCC1143 *GJB3*^*WT*^ cells to mammary adipocytes from all three patients (Fig. 3E). However, depletion of Cx31 expression by 1/3 or 2/3 in the *GJB3*^*Med*^ and *GJB3*^*Low*^ lines, respectively, resulted in a significant decrease in dye transfer compared to *GJB3*^*WT*^ control cells (Fig. 3E). These data suggest that functional gap junctions form between TN breast cancer cells and adipocytes in a Cx31-dependent manner.

To determine if breast cancer cell gap junctions are permeable to cAMP, we treated a panel of human TN and RP cell lines with CBX for 24 hours to inhibit pan-gap junction function and ascertain if cAMP was retained in the tumor cells. In 5 of 6 lines tested we found marked increases in the levels of intracellular cAMP concentration in CBX-versus vehicle-treated cells (Fig. 3F). Additionally, significantly higher concentrations of cAMP were observed in high MYC TN cells in comparison to low MYC TN or RP cells (Fig. 3F). The increase in intracellular cAMP upon pan-gap junction inhibition in 5 of 6 lines examined suggests that breast cancer cell gap junctions are indeed permeable to cAMP.

We next tested whether cAMP is directly transferred from breast cancer cells to adipocytes and if the abundance of Cx31 alters transfer. HCC1143 *GJB3*^*WT*^, *GJB3*^*Med*^ or *GJB3*^*Low*^ cells were seeded and loaded with a fluorescent cAMP analogue (fluo-cAMP). These monolayer cultures were then co-cultured in direct contact with primary mammary adipose tissue from three healthy individuals (WD47558, WD46812, WD50344), and incubated for 5 hours. Adipocytes were then isolated from the tumor cells and assayed for fluo-cAMP. We found that cAMP transfer occurred from control cells to adipocytes from all three patients (Fig. 3G). However, as we observed with transfer of gap junction-permeable dye (Fig. 3E), depletion of Cx31 resulted in a significant reduction of cAMP transfer (Fig. 3G). Thus, cAMP is transferred from TN breast cancer cells to adipocytes in a Cx31-dependent manner.

We next sought to determine if downstream cAMP signaling is activated in adipocytes in a tumor-adipocyte gap junction-dependent manner. To determine if cAMP signaling is activated in adipocytes upon cell-cell contact with breast cancer cells, we used a primary mouse preadipocyte model that can be differentiated to adipocytes *in vitro* [18, 38]. This model is ideal to study downstream signaling during co-culture because changes in adipocyte transcription can be assayed via qRT-PCR using murine-specific primers. Adipocytes were terminally differentiated and then HS578T and HCC1143 *GJB3* partial depletion cell lines were seeded directly on top of adipocyte cultures. After co-culturing the cells for 24 hours we extracted RNA and assayed for changes in murine-specific (thus adipocyte-specific in this system) expression of UCP1, a known cAMP-responsive gene in adipocytes [18], to measure cAMP signaling. We also assayed for mouse aP2 expression as a marker of adipocyte differentiation. Our positive control, forskolin, robustly induced UCP1 expression compared to vehicle-treated cells (Fig. 3H). The HCC1143 *GJB3*^*WT*^ and *GJB3*^*Med*^ lines both induced adipocyte UCP1 expression, but UCP1 induction was significantly reduced in the *GJB3*^*Low*^ co-cultures (Fig. 3H). In contrast, none of the HS578T lines, including the *GJB3*^*WT*^ control, were capable of inducing UCP1 expression (Fig. 3H). All conditions, including forskolin treatment, resulted in reduced aP2 expression (Fig. 3H), suggesting effects on adipocyte differentiation are distinct from those observed on cAMP signaling. Given that Cx31 expression is similar in HS578T *GJB3*^*WT*^ and HCC1143 *GJB3*^*Low*^ cells (Fig. 3C), and that neither activate cAMP signaling (Fig. 3H), it is possible that a Cx31 expression threshold is required for breast cancer cells to activate cAMP signaling in adjacent adipocytes. Although direct transfer of cAMP among adipocytes in a homotypic interaction has been described [21], this is the first description of gap junction-dependent activation of adipocyte lipolysis in a heterotypic manner, by a tumor cell.

Finally, we sought to determine the contribution of breast cancer Cx31-dependent gap junctions to tumorigenesis. We found that HS578T *GJB3*^*Med-1*^ and *GJB3*^*Med-2*^, and HCC1143 *GJB3*^*Med*^ cell lines did not display a difference in proliferation compared to their respective *GJB3*^*WT*^ control lines (Fig. 4A). In contrast, HCC1143 *GJB3*^*Low*^ cells demonstrate a significant reduction in proliferation, while maintaining 93.7% viability relative to Cas9 controls (Fig. 4A). These data suggest that, even in the absence of breast cancer cell-adipocyte interaction, Cx31 may promote breast cancer cell proliferation. To determine the contribution of Cx31 to breast tumorigenesis *in vivo*, we transplanted each of the HS578T and HCC1143 Cx31 partial depletion lines into mammary fat pads of immunocompromised NOD-SCID/gamma (NSG) female mice and assayed for time of tumor onset and ethical endpoint (when tumor reaches 2cm in any dimension). Remarkably, with the HS578T lines, in which partial *GJB3* knockout had no effect on cell proliferation *in vitro* (Fig. 4A), 0/10 mice that received HS578T *GJB3*^*Med-1*^ or *GJB3*^*Med-2*^ xenografts (5 per line) developed tumors within 180 days (Fig. 4B). Among the HCC1143 lines, the *GJB3*^*Med*^ line displayed a significant delay in both tumor onset and time to ethical endpoint, while only 3 of 5 mice transplanted with the *GJB3*^*Low*^ line developed tumors, and none reached ethical endpoint within 180 days (Fig. 4B). We performed an independent xenograft model wherein inducible Cx31 hairpins were transduced into the TN-MYC^High^ BT549 human breast cell line and found that Cx31 depletion significantly enhanced tumor-free survival compared to controls (Fig. 4C). Our data indicate that decreasing Cx31 expression is sufficient to impair tumor growth, suggesting that gap junctions promote breast tumorigenesis *in vivo*.

**Fig. 4.**
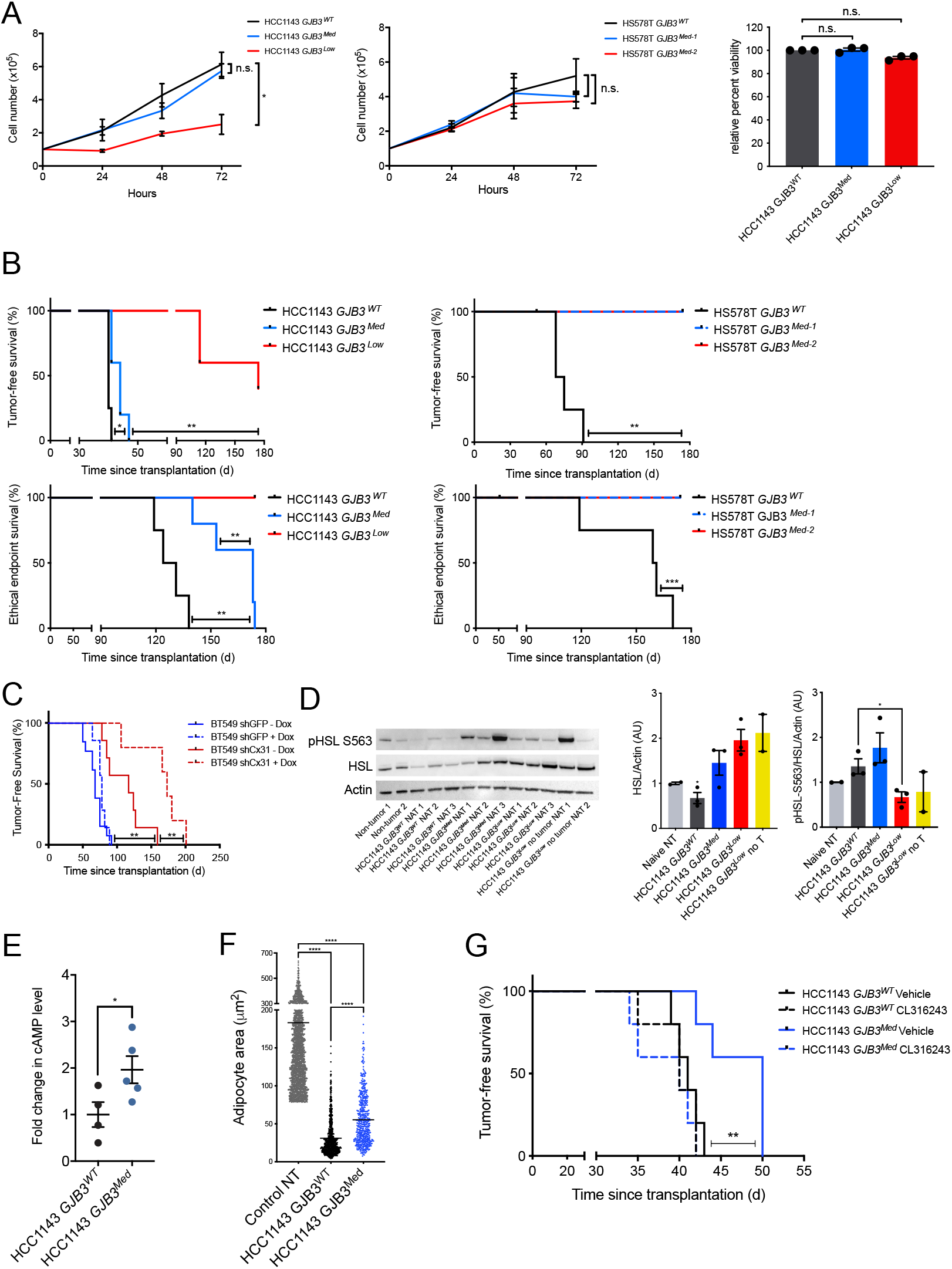
Cx31 loss impairs breast cancer cell growth in vitro, tumorigenesis, and activation of lipolysis in adjacent adipocytes in vivo. (A) Cell growth of indicated Cx31 partial expression loss lines in HCC1143 (left) and HS578T (middle) over 72 hours, and cell viability at 72 hours of indicated partial GJB3 depletion lines normalized to WT control (right). (B) Kaplan-Meier analysis of tumor onset (top) and ethical endpoint survival (bottom) of mice bearing indicated Cx31 partial expression loss orthotopic xenografts (n = 5 per group). Ethical endpoint survival indicates the percentage of mice bearing xenografts < 2cm in any dimension. (C) Kaplan-Meier analysis of tumor onset in mice bearing indicated xenografts with inducible Cx31 (shCx31) or GFP (shGFP) hairpin, with doxycycline (shGFP n = 7, shCx31 n = 5) and without doxycycline (shGFP n = 13, shCx31 n = 5). (D) Immunoblot analysis (left) showing expression levels of HSL and phosphorylated HSL (pHSL S563) in healthy non-tumor mammary gland and NAT from mice bearing indicated Cx31 partial expression loss xenografts or mice that were transplanted, but subsequently did not develop a tumor. Quantification of total HSL (middle) and of pHSL/HSL ratio (right), normalized to b-actin levels. (E) Fold change in cAMP levels in HCC1143 GJB3Med xenografts versus HCC1143 GJB3WT xenografts. (F) Adipocyte area adjacent to HCC1143 GJB3Med xenografts (n = 5) and HCC1143 GJB3WT xenografts (n = 4) and area in control non-tumor (NT) mice (n=3). The black line indicates mean adipocyte area. Each point represents an individual adipocyte. (G) Kaplan-Meier analysis of tumor onset of mice bearing indicated Cx31 partial expression loss orthotopic xenografts (n = 5 per group) and treated with vehicle or with 1mg/kg CL316243. For (D) and (E) mean ± s.e.m. is shown. *P < 0.05, **P < 0.01, ***P < 0.001, ****P < 0.0001; unpaired two-tailed t test (A) and (E), log-rank test (B), (C) and (G), ordinary one-way ANOVA (D) and (F).

We sought to clarify the effects of Cx31 on lipolysis versus other effects on tumor growth. To determine if control and Cx31 partial expression loss tumors differentially induced lipolysis, we collected tumor and NAT from HCC1143 *GJB3*^*WT*^, *GJB3*^*Med*^ and *GJB3*^*Low*^ tumor-bearing mice, as well as residual mammary glands from the two *GJB3*^*Low*^ mice that were transplanted, but never developed tumors. Using immunoblot analysis, we probed for markers of lipolysis. Notably, a marked reduction in total HSL expression was found in 3 of 3 HCC1143 *GJB3*^*WT*^ NAT samples compared to control tissues (Fig. 4D), consistent with persistent activation of lipolysis leading to HSL downregulation [23, 24]. In contrast, we did not observe a consistent change in HSL expression in any of the other NAT samples analyzed from tumors with partial Cx31 expression loss (Fig. 4D). Interestingly, we found a marked increase in phospho-HSL/HSL ratio in both the HCC1143 *GJB3*^*WT*^ and *GJB3*^*Med*^ NAT samples, but this difference was significantly reduced in HCC1143 *GJB3*^*Low*^ NAT (Fig. 4D). The increase in phospho-HSL/HSL in *GJB3*^*Med*^ NAT may be due to alternative modes of lipolysis activation, such as secreted pro-lipolytic cytokines [4], which is congruent with the observed increase in UCP1 expression during *GJB3*^*Med*^-adipocyte co-culture (Fig. 3H). To further interrogate lipolytic signaling in NAT, we probed for cAMP abundance in HCC1143 *GJB3*^*WT*^ and *GJB3*^*Med*^ tumors by mass spectrometry. We found a significant increase in intratumoral cAMP level in HCC1143 *GJB3*^*Med*^ tumors compared to the *GJB3*^*WT*^ control tumors (Fig. 4E), consistent with diminished transfer of cAMP to NAT. We examined *GJB3*^*WT*^ and *GJB3*^*Med*^ tumors and associated NAT, and assayed for differences in adjacent adipocyte size, as an indicator of lipolysis. We found a significant increase in the average size of adipocytes adjacent to *GJB3*^*Med*^ tumors compared to *GJB3*^*WT*^ control tumors (Fig. 4F), again supporting a decreased induction of lipolysis in NAT from Cx31 partial knockout tumors.

Finally, if the delay in HCC1143 *GJB3*^*Med*^ tumor onset (Fig. 4B) was due to an inability to activate lipolysis in adjacent adipocytes, we reasoned that pharmacological activation of lipolysis should rescue this phenotype. Indeed, we found that daily intra-peritoneal injection of CL316243, a specific β3-receptor agonist known to activate lipolysis *in vivo* [39], completely rescued the delay in tumor onset observed in HCC1143 *GJB3*^*Med*^ tumors, but did not further promote the growth of HCC1143 *GJB3*^*WT*^ tumors (Fig. 4G). Taken together, these data indicate that cAMP signaling, and lipolysis are activated in breast tumor-adjacent adipocytes in a Cx31-dependent manner *in vivo*.

In summary, we find that lipolysis is activated in breast cancer-adjacent adipose tissue and that functional gap junctions form between breast cancer cells, and between breast cancer cells and adipocytes. In addition, cAMP is transferred via breast cancer cell gap junctions, and cAMP signaling is activated in adipocytes adjacent to breast cancer cells in a gap junction-dependent manner. Finally, we established a previously unappreciated, functional role for Cx31-dependent gap junctions in promoting breast tumor growth and activation of lipolysis in tumor-adjacent adipose tissue *in vivo*, which may represent a new therapeutic target to treat pro-lipolytic breast tumors. Furthermore, the recent discovery of gap junction formation and pro-tumorigenic signal exchange between brain metastatic carcinoma cells and astrocytes [30] suggests that gap junction-dependent heterotypic interaction between tumor and non-tumor cells may be an emerging hallmark of tumorigenesis.

## Acknowledgements

This work was supported, in part, by the US Department of Defense–Congressionally Directed Medical Research Programs’ Era of Hope Scholar award W81XWH-12-1-0272 (A.G.), the US National Institutes of Health (NIH) grants R01 CA 0447 (A.G.) and R01 CA056721 (Z.W.), the Atwater Foundation (A.G.), the Breast Cancer Research Fund (H.R.), the EMBO postdoctoral fellowship ALTF 159-2017 (Ju.W.), the US K99/R00 NIH Pathway to Independence Award DK110426 (K.S.), and the US NIH F99/K00 Predoctoral to Postdoctoral Transition Award F99CA212488 (R.C.) and the US NIH F31 Predoctoral Individual National Research Service Award F31CA243468 (J.W.). The authors thank A. Welm for guidance in the use of patient-derived xenografts, A. Tward for technical guidance and helpful discussions, K.A. Fontaine for helpful discussions and comments on the manuscript, and S. Samson for a helpful consumer advocate perspective and guidance on our work. R.C. and A.G. conceived of research. J.W. and R.C. designed and contributed to all *in vitro* and *ex vivo* studies, and to the *in vivo* mouse studies, contributed to all data analysis, and wrote the manuscript. S.M. analyzed 3CB data. L.J.Z. designed LCM and conducted proteomic analyses. S.M. conducted the LCM. D.A. conducted lipolysis signature enrichment and scRNA-seq analyses. A.B. contributed to immunofluorescent staining and microscopy, and generation of Cx31 partial knockout lines. D.V. characterized Cx31 partial depletion lines, generated Cx31 inducible short hairpin lines, contributed to Cx31 knockdown and CL316243 mouse studies and provided helpful discussions. S.K.A. contributed to generation of Cx31 inducible short hairpin lines. R.N. contributed to co-culture studies, immunofluorescent staining, tissue clearing and microscopy, and provided valuable discussions. Y.C. conducted preadipocyte differentiation. C.B. and S.L. conducted mass spectrometry for cAMP. C.M. generated Cas9-expressing cancer cell lines. Ju.W. and E.W. isolated primary mammary adipose tissue. J.D.G. and D.S. conducted FACS analysis. K.S. provided cAMP-dependent lipolysis signature. K.M.A. supervised FACS analysis. Z.W. supervised primary mammary adipose tissue isolation and provided valuable discussion. D.K.N. supervised mass spectrometry for cAMP and provided valuable discussion. S.K. supervised preadipocyte and cAMP-dependent lipolysis studies and provided valuable discussion. A.J.B supervised enrichment analysis. M.E.S. and D.C.L. designed and supervised LCM and proteomics. M.G. contributed to RNA-seq analysis. H.N. contributed to co-culture studies. H.R. provided valuable discussion. G.K. designed and conducted histological analyses and provided valuable discussion. J.A.S. supervised the 3CB study and data analysis, and provided valuable discussion. A.G. supervised the study, and provided valuable discussion and intellectual input. All authors edited the manuscript. All data and code related to these studies are available in the main text, supplementary materials and indicated repositories. The raw RNAseq data will be deposited on GEO. The authors declare no competing interest.

---

**Fig. S1.**
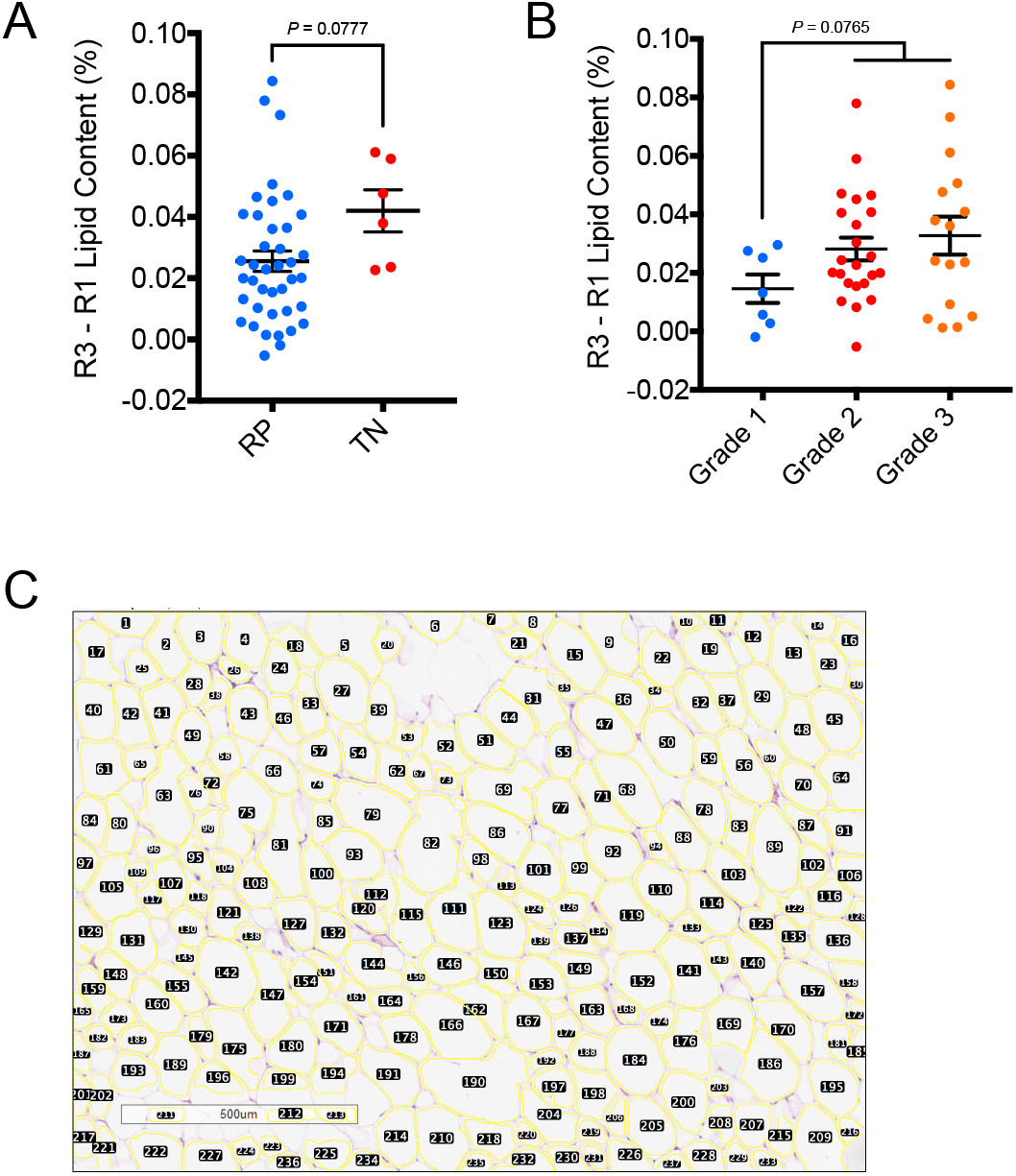
NAT lipid content by receptor status and tumor grade, and adipocyte area quantification. (A) Change in lipid content in R3 of NAT versus R1 of NAT from TN and RP patients. (B) Change in lipid content in R3 of NAT versus R1 of NAT from grade 1, 2 and 3 patients. (C) Example of Adiposoft software output on manual mode before curation to identify whole, individual adipocytes. *P* values indicated; unpaired two-tailed *t* test (A) and (B).

**Fig. S2.**
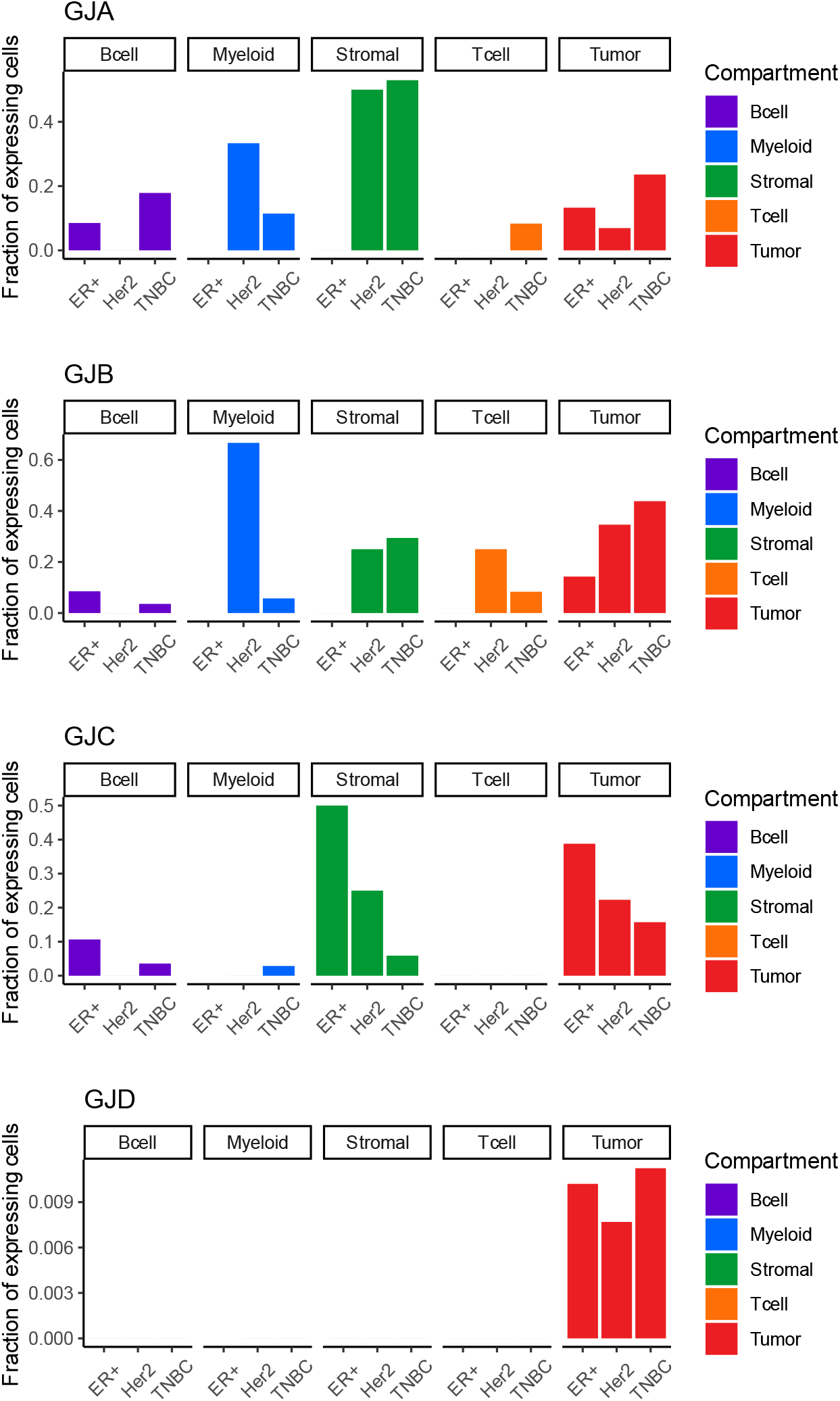
Fraction of cells expressing gap junction family by tumor compartment cell type. Single cell (n = 515) RNA-seq of B cell (n = 83), myeloid cell (n = 38), stromal cell (n = 23), T cell (n = 54) and tumor (n = 317) cell compartments from the patient (n = 11) tumor microenvironment

**Fig. S3.**
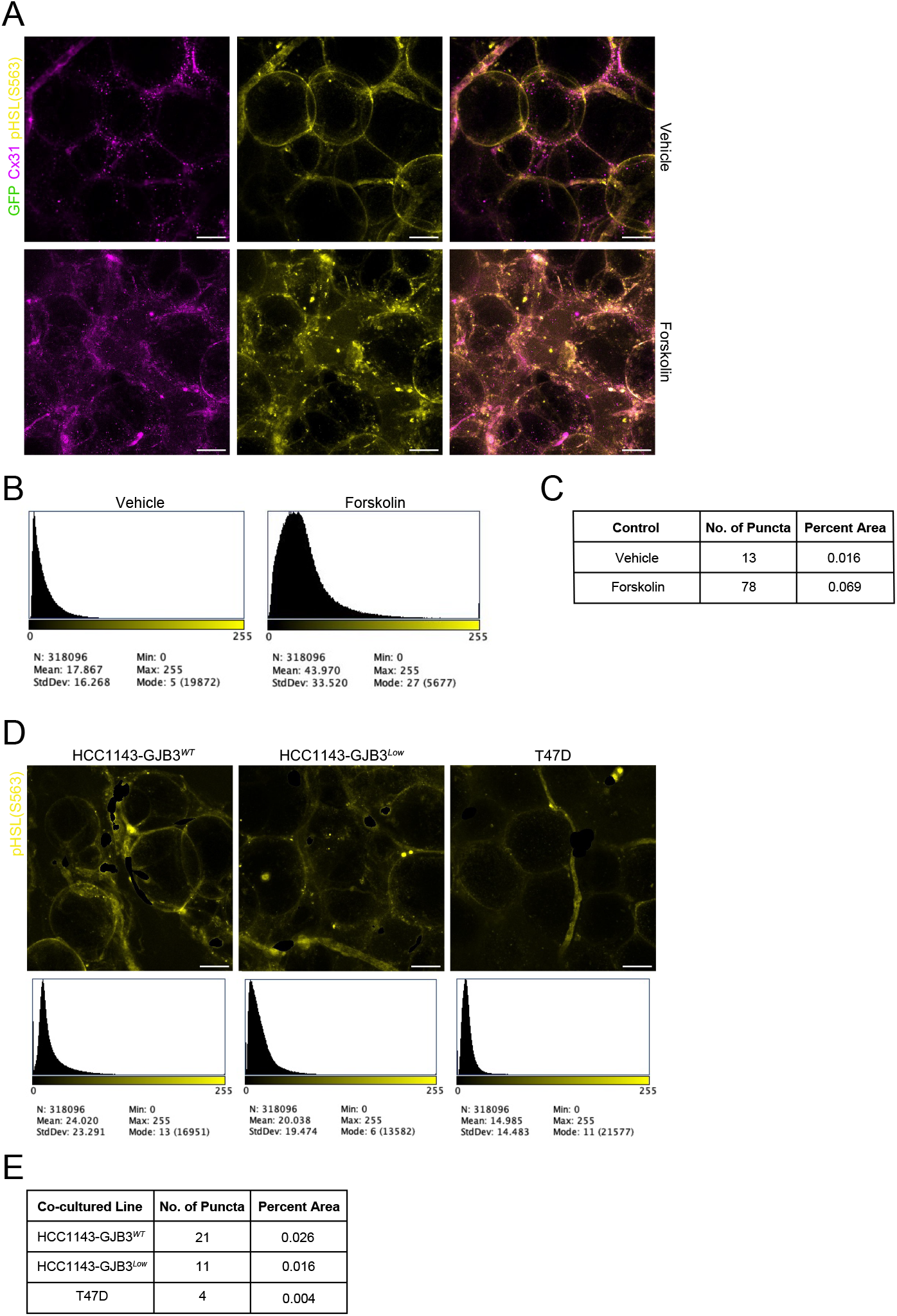
Quantification of lipolysis signaling in primary mammary adipose tissue from indicated cancer cell co-cultures and controls. (A) Staining with Cx31 (magenta) and pHSL(S563) (yellow) of control mammary adipose tissue injected with either vehicle (top) or 10 μM forskolin (bottom) and cultured for 24 hours. Scale bar, 50 μm. (B) Histogram of pHSL(S563) expression in indicated coculture control tissues. (C) Quantification of pHSL(S563) puncta number and percent total area in indicated co- culture control tissues. (D) Staining (top) and histogram (bottom) of pHSL(S563) in mammary adipose tissue co-cultured with indicated GFP-tagged cancer cell lines. Cancer cell pHSL(S563) signal was masked out using GFP tag. Scale bar, 50 μm. (E) Quantification of pHSL(S563) puncta number and percent total area in mammary adipose tissue co-cultured with indicated cell line.

**Fig. S4.**
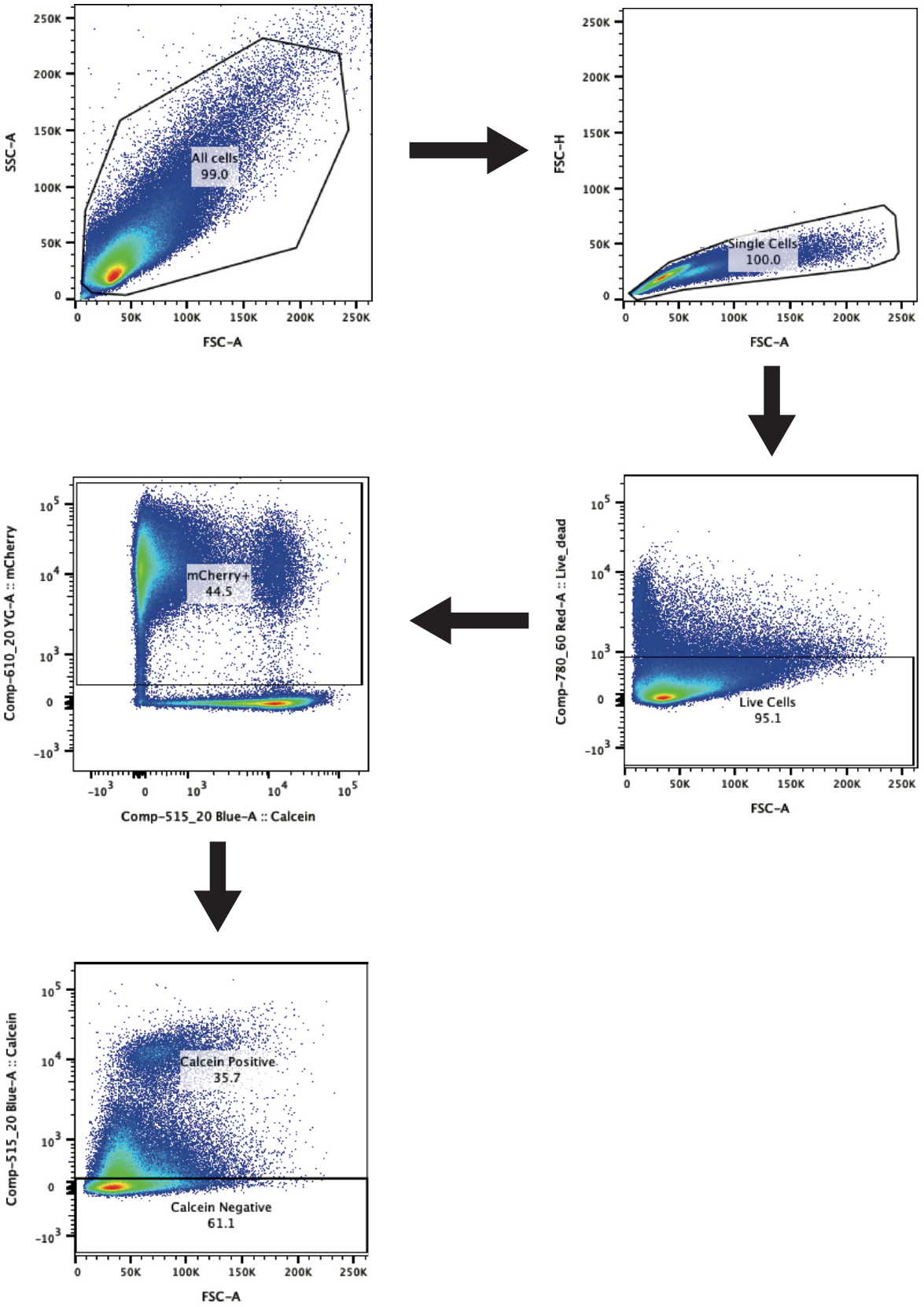
Flow cytometry gating strategy to identify mCherry-positive, CalceinAM-positive cells. Representative experimental control sample, HCC1143 cells. Side scatter and forward scatter were used to distinguish all cells from debris. Forward scatter was used to distinguish singlets (single cells) from all cells. Live cells were identified as negative for live/dead staining. Live single cells positive for mCherry were identified. Of mCherry-positive cells, CalceinAM-positive and -negative populations were distinguished.

## Supplementary Table Legends

Table S1. Patient ID, receptor status, histological section availability, percent lipid content (lipid content / lipid + water + protein content) of L, R1, R2 and R3, and Scarff-Bloom-Richardson (SBR) grade from patients (*n* = 46) with invasive breast cancer.

Table S2. LC-MS/MS of LCM samples from 75 patients with invasive breast cancer and 42 healthy subjects. (A) Sample number, ID number, tissue type, and tumor subtype (when applicable) of 75 patients and 42 healthy subjects. (B) Spectral counts of proteins detected via LC-MS/MS from samples in (A).

Table S3. RNA expression changes in MTB-TOM tumors (*n* = 11) compared to non-tumor mammary glands (*n* = 3). Differential expression analysis was performed using the ‘limma’ R package [49].

**Table S4.**
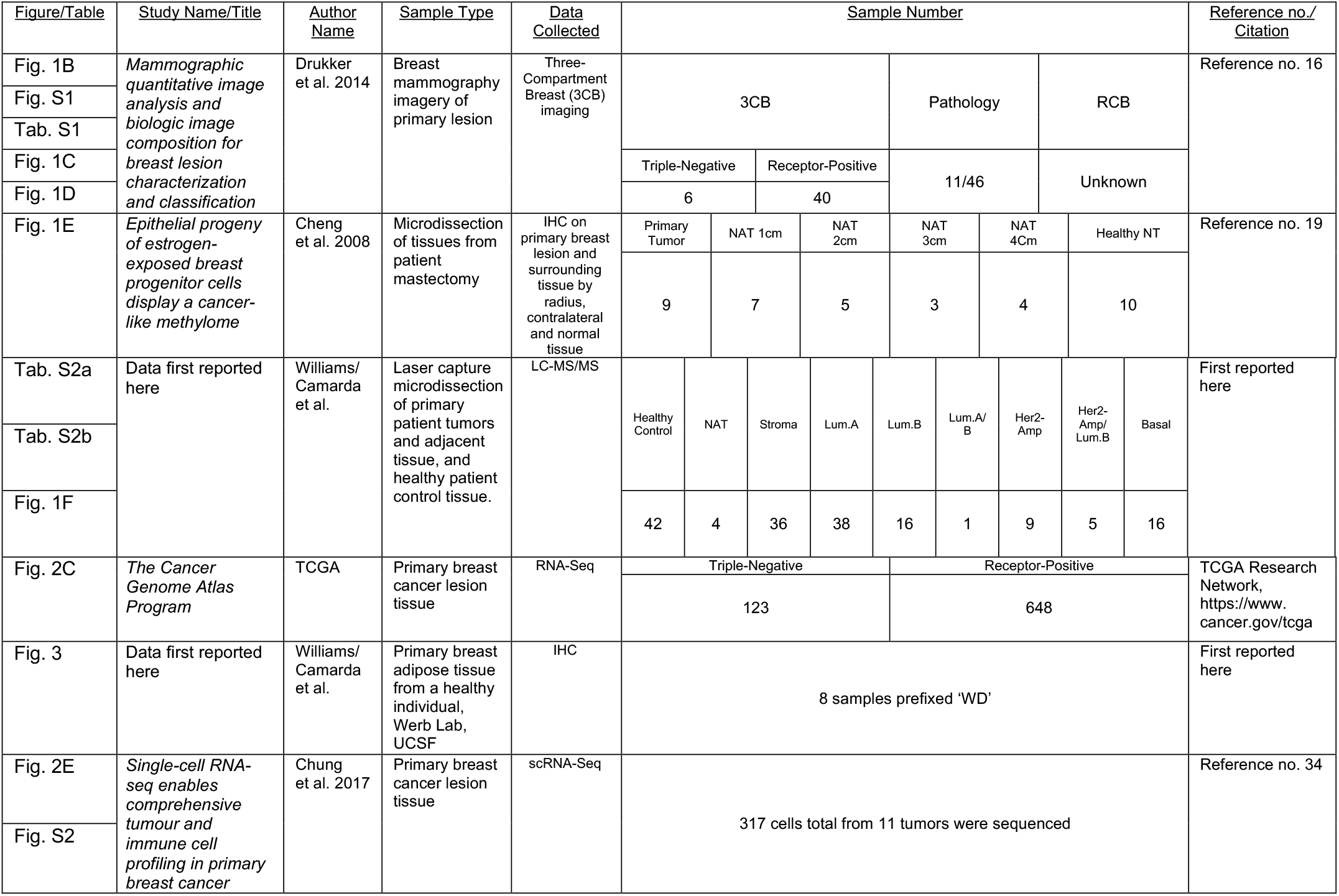
Sample specifications and study sources for applied clinical samples and datasets.

